# Transmission of SARS-COV-2 from China to Europe and West Africa: a detailed phylogenetic analysis

**DOI:** 10.1101/2020.10.02.323519

**Authors:** Wasco Wruck, James Adjaye

## Abstract

**Background:** SARS-CoV-2, the virus causing the Covid-19 pandemic emerged in December 2019 in China and raised fears that it could overwhelm healthcare systems worldwide. In June 2020, all African countries registered human infections with SARS-CoV-2.

The virus is mutating steadily and this is monitored by a well curated database of viral nucleotide sequences from samples taken from infected individual thus enabling phylogenetic analysis and phenotypic associations.

**Methods:** We downloaded from the GISAID database, SARS-CoV-2 sequences established from four West African countries Ghana, Gambia, Senegal and Nigeria and then performed phylogenetic analysis employing the nextstrain pipeline. Based on mutations found within the sequences we calculated and visualized statistics characterizing clades according to the GISAID nomenclature.

**Results:** We found country-specific patterns of viral clades: the later Europe-associated G-clades predominantly in Senegal and Gambia, and combinations of the earlier (L, S, V) and later clades in Ghana and Nigeria. Contrary to our expectations, the later Europe-associated G-clades emerged before the earlier clades. Detailed analysis of distinct samples showed that some of the earlier clades might have circulated latently and some reflect migration routes via Mali and Tunisia.

**Conclusions:** The distinct patterns of viral clades in the West African countries point at its emergence from Europe and China via Asia and Europe. The observation that the later clades emerged before the earlier clades could be simply due to founder effects or due to latent circulation of the earlier clades. Only a marginal correlation of the G-clades associated with the D614G mutation could be identified with the relatively low case fatality (0.6-3.2).

**Key messages:**

- Ghana and Nigeria have a combination of earlier (L, V, S) and later Europe-associated G-clades of SARS-CoV-2, therefore pointing to multiple introductions while in Senegal and Gambia Europe-associated G-clades predominate pointing to introductions mainly from Europe.
- Surprisingly, the later G-clades emerged before the earlier clades (L, V, S)
- Detailed phylogenetic analysis points at latent circulation of earlier clades before the first registered cases.
- Phylogenetic analysis of some cases points at migration routes to Europe via Tunisia, Egypt and Mali.
- A marginal correlation of r=0.28 between the percentage of the D614G mutation defining the G-clades and case-fatality can be detected.

## Introduction

The Covid-19 pandemic resulting from the SARS-CoV-2 coronavirus infection which emerged in December 2019 in Wuhan, China, spread all over the world and after a delay of a few months also appeared on the African continent. Early in June 2020, all African countries registered human infections with SARS-CoV-2. Starting with the first sequenced human sample of SARS-CoV-2, several mutations of the virus sequence arose which could be grouped into clades allowing associations with regional prevalences. In this study, we focus on samples from West Africa which was the region from which the first African SARS-CoV-2 sequences became available. We aimed at analysing phylogenetic characteristics possibly giving clues about the distribution between countries and eventually even about putative severity changes between specific clades. We use the nomenclature of clades suggested by the GISAID initiative and adopted in several publications. Previous studies reported potential impact of the D614G amino acid mutation which is induced by the A23403G single nucleotide polymorphism (SNP) (1) and associated with the branch of the phylogenetic tree referred to as clade G.

Brufsky hypothesized that the higher number of deaths on the East coast of the United States compared to the West coast could be due to the higher prevalence of the D614G mutation on the East coast (2). The D614G mutation has been suggested to affect the adherence of the virus to the cell membrane and as a consequence results in higher virulence. Supportive evidence was reported in mice (3), (4). Korber et al. hypothesized that the D614 amino acid on the surface of the spike protein protomer region S1 of the virus could have a hydrogen bond to the T859 amino acid in the S2 region residing on the membrane (1). Furthermore, they showed that clade G rapidly starts to replace other clades associated with the D614 amino acid in each country entered (1). In mid-March 2020, the G clade was found almost exclusively in Europe (5) but soon after spread all over the world. Korber et al. see its origin from China or Europe (1). In China, four early samples carried the D614G mutation. One sample from January 24^th^ 2020 had only the A23403G (D614G) but not the C3037T and C14408T mutations which usually associate with A23403T in clade G. Three samples with the D614G were related to the first German sample. In Europe the first German sample from January 28^th^ carried the A-to-G mutation at nucleotide position 23403 (D614G) mutation and the C-to-T mutation at position 3037, but not the mutation at position 14408. The first sample carrying all of the above mentioned mutations plus the C241T in the Untranslated Region (UTR) was identified in Italy on February 20^th^ 2020 (1). Another interesting feature of the G clades is that the associated C14408T mutation adjacent to the RNA dependent RNA polymerase (RdRp) putatively increases the mutation rate as Pachetti *et al*. report (5).

Detailed analyses of virus evolution have been performed for some countries, e.g. France (6), New York (7) and India (8). For France it could be deduced by the distinction between clade G and the earlier phylogenetic branches that the first SARS-CoV-2 did not lead to local transmission while the clade G was circulating for a considerable time before the first recorded case which was of clade G and had no travel events or traveller contact (6).

Since the declaration of Covid-19 as pandemic by the WHO on March 11^th^ 2020, fears were expressed that it could overwhelm weaker healthcare systems as existing in many African countries. Furthermore, hygiene, social distancing and lockdown face many challenges in countries with high percentages without clean running water, cramped confines and the dependence on a daily income. However, Africa has the advantage of a very young population, e.g. in Sub-Saharan Africa with a median age of 19.7 years (9) for which in average milder etiopathologies can be expected. Additionally, for the early outbreak a study evaluating air traffic from affected regions in China calculated relatively low transmission risks for most African countries except South Africa and Ethiopia (10). For later phases, Cabore et al. proposed a model estimating risk of exposure for African countries based on a Hidden Markov model which accounts for factors such as gathering, weather, distribution and hygiene with the conclusion that with respect to the high calculated infection rates effective containment is indispensable (11).

Here, we analysed SARS-CoV-2 nucleotide sequenes from samples obtained from the West African countries of Gambia, Ghana, Nigeria and Senegal in order to identify characteristic mutations and to dissect their patterns of distribution.

## Methods

### Sample collection

We downloaded SARS-CoV-2 viral sequences for West African samples and reference samples from European, North and South American countries and China from the GISAID database of June 2020. The samples used in this study are shown in Table 1.

**Table 1:**
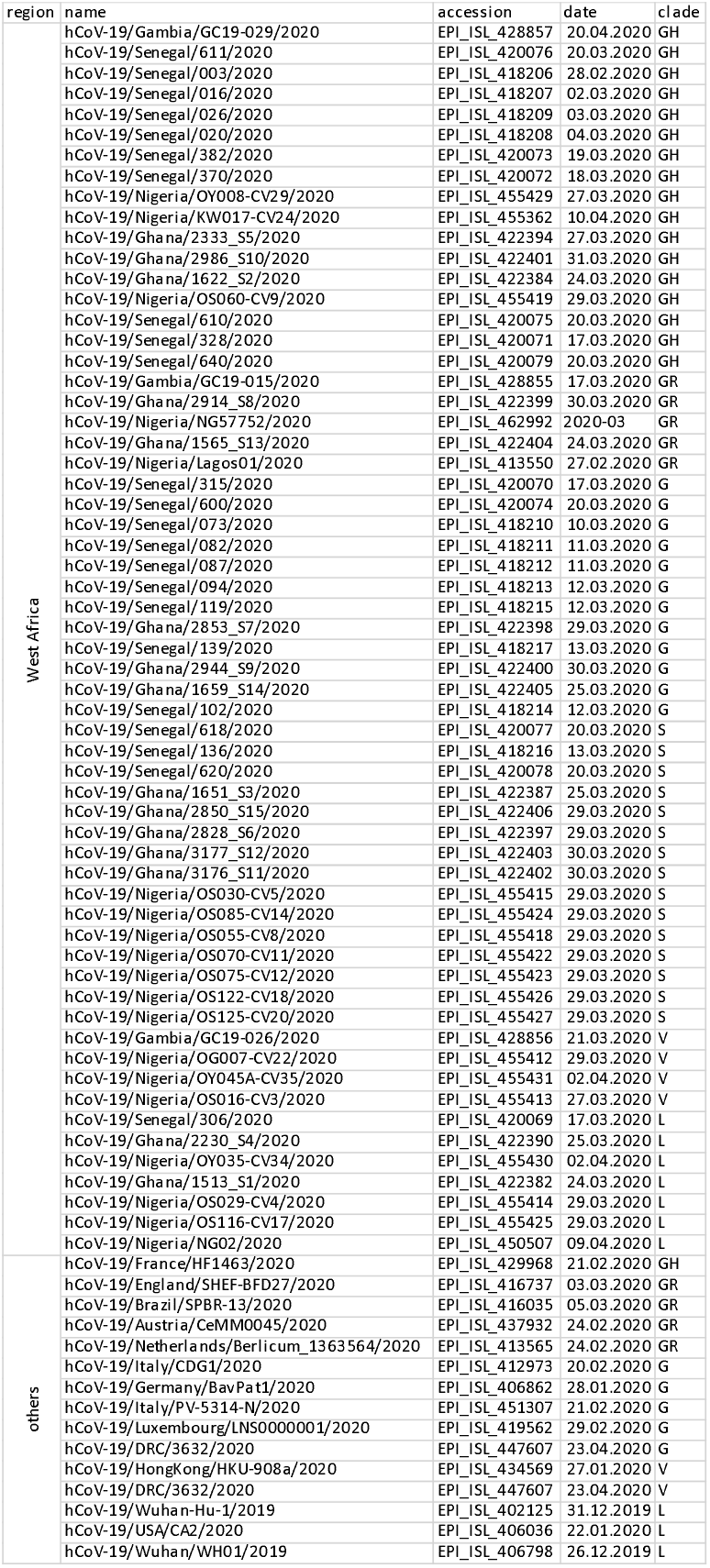
SARS-CoV-2samples used for the phylogenetic analysis

| region | name | accession | date | clade |
| --- | --- | --- | --- | --- |
| West Africa | hCoV-19/Gambia/GC19-029/2020 | EPI_ISL_428857 | 20.04.2020 | GH |
|  | hCoV-19/Senegal/611/2020 | EPI_ISL_420076 | 20.03.2020 | GH |
|  | hCoV-19/Senegal/003/2020 | EPI_ISL_418206 | 28.02.2020 | GH |
|  | hCoV-19/Senegal/016/2020 | EPI_ISL_418207 | 02.03.2020 | GH |
|  | hCoV-19/Senegal/026/2020 | EPI_ISL_418209 | 03.03.2020 | GH |
|  | hCoV-19/Senegal/020/2020 | EPI_ISL_418208 | 04.03.2020 | GH |
|  | hCoV-19/Senegal/382/2020 | EPI_ISL_420073 | 19.03.2020 | GH |
|  | hCoV-19/Senegal/370/2020 | EPI_ISL_420072 | 18.03.2020 | GH |
|  | hCoV-19/Nigeria/OY008-CV29/2020 | EPI_ISL_455429 | 27.03.2020 | GH |
|  | hCoV-19/Nigeria/KW017-CV24/2020 | EPI_ISL_455362 | 10.04.2020 | GH |
|  | hCoV-19/Ghana/2333_S5/2020 | EPI_ISL_422394 | 27.03.2020 | GH |
|  | hCoV-19/Ghana/2986_S10/2020 | EPI_ISL_422401 | 31.03.2020 | GH |
|  | hCoV-19/Ghana/1622_S2/2020 | EPI_ISL_422384 | 24.03.2020 | GH |
|  | hCoV-19/Nigeria/OS060-CV9/2020 | EPI_ISL_455419 | 29.03.2020 | GH |
|  | hCoV-19/Senegal/610/2020 | EPI_ISL_420075 | 20.03.2020 | GH |
|  | hCoV-19/Senegal/328/2020 | EPI_ISL_420071 | 17.03.2020 | GH |
|  | hCoV-19/Senegal/640/2020 | EPI_ISL_420079 | 20.03.2020 | GH |
|  | hCoV-19/Gambia/GC19-015/2020 | EPI_ISL_428855 | 17.03.2020 | GR |
|  | hCoV-19/Ghana/2914_S8/2020 | EPI_ISL_422399 | 30.03.2020 | GR |
|  | hCoV-19/Nigeria/NG57752/2020 | EPI_ISL_462992 | 2020-03 | GR |
|  | hCoV-19/Ghana/1565_S13/2020 | EPI_ISL_422404 | 24.03.2020 | GR |
|  | hCoV-19/Nigeria/Lagos01/2020 | EPI_ISL_413550 | 27.02.2020 | GR |
|  | hCoV-19/Senegal/315/2020 | EPI_ISL_420070 | 17.03.2020 | G |
|  | hCoV-19/Senegal/600/2020 | EPI_ISL_420074 | 20.03.2020 | G |
|  | hCoV-19/Senegal/073/2020 | EPI_ISL_418210 | 10.03.2020 | G |
|  | hCoV-19/Senegal/082/2020 | EPI_ISL_418211 | 11.03.2020 | G |
|  | hCoV-19/Senegal/087/2020 | EPI_ISL_418212 | 11.03.2020 | G |
|  | hCoV-19/Senegal/094/2020 | EPI_ISL_418213 | 12.03.2020 | G |
|  | hCoV-19/Senegal/119/2020 | EPI_ISL_418215 | 12.03.2020 | G |
|  | hCoV-19/Ghana/2853_S7/2020 | EPI_ISL_422398 | 29.03.2020 | G |
|  | hCoV-19/Senegal/139/2020 | EPI_ISL_418217 | 13.03.2020 | G |
|  | hCoV-19/Ghana/2944_S9/2020 | EPI_ISL_422400 | 30.03.2020 | G |
|  | hCoV-19/Ghana/1659_S14/2020 | EPI_ISL_422405 | 25.03.2020 | G |
|  | hCoV-19/Senegal/102/2020 | EPI_ISL_418214 | 12.03.2020 | G |
|  | hCoV-19/Senegal/618/2020 | EPI_ISL_420077 | 20.03.2020 | S |
|  | hCoV-19/Senegal/136/2020 | EPI_ISL_418216 | 13.03.2020 | S |
|  | hCoV-19/Senegal/620/2020 | EPI_ISL_420078 | 20.03.2020 | S |
|  | hCoV-19/Ghana/1651_S3/2020 | EPI_ISL_422387 | 25.03.2020 | S |
|  | hCoV-19/Ghana/2850_S15/2020 | EPI_ISL_422406 | 29.03.2020 | S |
|  | hCoV-19/Ghana/2828_S6/2020 | EPI_ISL_422397 | 29.03.2020 | S |
|  | hCoV-19/Ghana/3177_S12/2020 | EPI_ISL_422403 | 30.03.2020 | S |
|  | hCoV-19/Ghana/3176_S11/2020 | EPI_ISL_422402 | 30.03.2020 | S |
|  | hCoV-19/Nigeria/OS030-CV5/2020 | EPI_ISL_455415 | 29.03.2020 | S |
|  | hCoV-19/Nigeria/OS085-CV14/2020 | EPI_ISL_455424 | 29.03.2020 | S |
|  | hCoV-19/Nigeria/OS055-CV8/2020 | EPI_ISL_455418 | 29.03.2020 | S |
|  | hCoV-19/Nigeria/OS070-CV11/2020 | EPI_ISL_455422 | 29.03.2020 | S |
|  | hCoV-19/Nigeria/OS075-CV12/2020 | EPI_ISL_455423 | 29.03.2020 | S |
|  | hCoV-19/Nigeria/OS122-CV18/2020 | EPI_ISL_455426 | 29.03.2020 | S |
|  | hCoV-19/Nigeria/OS125-CV20/2020 | EPI_ISL_455427 | 29.03.2020 | S |
|  | hCoV-19/Gambia/GC19-026/2020 | EPI_ISL_428856 | 21.03.2020 | V |
|  | hCoV-19/Nigeria/OG007-CV22/2020 | EPI_ISL_455412 | 29.03.2020 | V |
|  | hCoV-19/Nigeria/OY045A-CV35/2020 | EPI_ISL_455431 | 02.04.2020 | V |
|  | hCoV-19/Nigeria/OS016-CV3/2020 | EPI_ISL_455413 | 27.03.2020 | V |
|  | hCoV-19/Senegal/306/2020 | EPI_ISL_420069 | 17.03.2020 | L |
|  | hCoV-19/Ghana/2230_S4/2020 | EPI_ISL_422390 | 25.03.2020 | L |
|  | hCoV-19/Nigeria/OY035-CV34/2020 | EPI_ISL_455430 | 02.04.2020 | L |
|  | hCoV-19/Ghana/1513_S1/2020 | EPI_ISL_422382 | 24.03.2020 | L |
|  | hCoV-19/Nigeria/OS029-CV4/2020 | EPI_ISL_455414 | 29.03.2020 | L |
|  | hCoV-19/Nigeria/OS116-CV17/2020 | EPI_ISL_455425 | 29.03.2020 | L |
|  | hCoV-19/Nigeria/NG02/2020 | EPI_ISL_450507 | 09.04.2020 | L |
| others | hCoV-19/France/HF1463/2020 | EPI_ISL_429968 | 21.02.2020 | GH |
|  | hCoV-19/England/SHEF-BFD27/2020 | EPI_ISL_416737 | 03.03.2020 | GR |
|  | hCoV-19/Brazil/SPBR-13/2020 | EPI_ISL_416035 | 05.03.2020 | GR |
|  | hCoV-19/Austria/CeMM0045/2020 | EPI_ISL_437932 | 24.02.2020 | GR |
|  | hCoV-19/Netherlands/Berlicum_1363564/2020 | EPI_ISL_413565 | 24.02.2020 | GR |
|  | hCoV-19/Italy/CDG1/2020 | EPI_ISL_412973 | 20.02.2020 | G |
|  | hCoV-19/Germany/BavPat1/2020 | EPI_ISL_406862 | 28.01.2020 | G |
|  | hCoV-19/Italy/PV-5314-N/2020 | EPI_ISL_451307 | 21.02.2020 | G |
|  | hCoV-19/Luxembourg/LNS0000001/2020 | EPI_ISL_419562 | 29.02.2020 | G |
|  | hCoV-19/DRC/3632/2020 | EPI_ISL_447607 | 23.04.2020 | G |
|  | hCoV-19/HongKong/HKU-908a/2020 | EPI_ISL_434569 | 27.01.2020 | V |
|  | hCoV-19/DRC/3632/2020 | EPI_ISL_447607 | 23.04.2020 | V |
|  | hCoV-19/Wuhan-Hu-1/2019 | EPI_ISL_402125 | 31.12.2019 | L |
|  | hCoV-19/USA/CA2/2020 | EPI_ISL_406036 | 22.01.2020 | L |
|  | hCoV-19/Wuhan/WHO1/2019 | EPI_ISL_406798 | 26.12.2019 | L |

### Construction of the phylogenetic tree

The phylogenetic tree was constructed using a pipeline adapted from the Zika virus pipeline on the nextstrain.org web page (12) employing the Augur (12), the MAFFT (13) and the IQ-tree (14) software. Details of steps which were performed:

First all West African and reference sequences in FASTA format were aligned employing the augur command:

> *augur align --sequences westafrica.fasta --reference-sequence sars_cov2_referencesequence.gb --output wa_aligned.fasta --fill-gaps*

which called MAFFT (13) with the command:

> *mafft --reorder --anysymbol --nomemsave --adjustdirection --thread 1 wa_aligned.fasta.to_align.fasta 1> wa_aligned.fasta 2> wa_aligned.fasta.log*

Metadata was extracted from the sequences FASTA via the augur command:

> *augur parse --sequences wa_aligned.fasta --fields strain accession date --output-sequences wa_aligned_parsed.fasta --output-metadata metadata.tsv*

Then a tree was built via the augur command:

> *augur tree --alignment wa_aligned_parsed.fasta --output wa_tree_raw.nwk*

Calling the IQ-tree algorithm (14) via this command:

> *iqtree -ninit 2 -n 2 -me 0.05 -nt 1 -s wa_aligned_parsed-delim.fasta -m GTR > wa_aligned_parsed-delim.iqtree.log*

The tree was refined via the Augur software calling TreeTime (15) for Maximum-Likelihood analysis inferring a time resolved phylogeny tree:

> *augur refine --tree wa_tree_raw.nwk --alignment wa_aligned_parsed.fasta --metadata metadata.tsv --output-tree wa_tree.nwk --output-node-data wa_branch_lengths.json --timetree --coalescent opt --date-confidence --date-inference marginal --clock-filter-iqd 4 --keep-polytomies*

Here, the command from the Zika pieline was adapted to --keep-polytomies to keep all samples. Metadata information was manually supplemented with country and region information and associated with the tree via a call to Augur:

> *augur traits --tree wa_tree.nwk --metadata metadata_countries.tsv --output wa_traits.json --columns region country --confidence*

Augur was called to infer ancestral states of discrete character again using TreeTime (15):

> *augur ancestral --tree wa_tree.nwk --alignment wa_aligned_parsed.fasta --output-node-data wa_nt_muts.json --inference joint*

Amino acid mutations were identified with the augur translate command:

> *augur translate --tree wa_tree.nwk --ancestral-sequences wa_nt_muts.json --reference-sequence sars_cov2_referencesequence.gb --output wa_aa_muts.json*

Results were exported via the augur command:

> *augur export v2 --tree wa_tree.nwk --metadata metadata_countries.tsv --node-data wa_branch_lengths.json wa_traits.json wa_nt_muts.json wa_aa_muts.json --colors colors.tsv --lat-longs lat_longs.tsv --auspice-config auspice_config.json --output wa_cov19.json*

### Visualization of the phylogenetic tree and annotation with mutations and clades

The phylogenetic tree was annotated with crucial mutations using the tool FigTree version 1.4.4 (http://tree.bio.ed.ac.uk/software/figtree/). Branches of the tree corresponding to clades following the nomenclature of GISAID were coloured distinctly.

### World map chart

The world map chart was built using the R-package Rworldmap (16). Clade distribution pie charts were copied to the distinct country locations. Connections between countries were based on the nextstrain Africa analysis and our own auspice analysis. Further connections between countries were retrieved from literature on virus introductions into countries or regions. The first patient on the West Coast of the United States returned from a journey to Wuhan, China (17). The first introductions in New York came from multiple independent infected individuals mainly from Europe (7). The first cases in France and Europe were Chinese travellers from the predominantly affected Hubei province who entered the county in mid-January and were tested positive on January 24^th^ 2020 (18). Patient zero in Germany was a Chinese resident from Wuhan visiting Germany (19). The Italian outbreak started with two Chinese travellers who arrived in Milan-Lombardy, went to Rome later on and were tested positive on January 31^st^ 2020 (20). The first Italian citizen was confirmed for Covid-19 on February 21^st^ 2020 in Lombardy (20). In the Netherlands, the first patient diagnosed on February 27^th^ 2020 had probably infected himself on a trip to Northern Italy between February 18^th^ and 21^st^ (21). The first cases in the UK returned from travels to the Chinese Hubei province and were tested positive for SARS-CoV-2 on January 30^th^ 2020 (22).

## Results

### Phylogenetic tree and diversity

The phylogenetic tree shown in Figure 1a displays similarities of West-African virus sequences with representative reference sequences from China and multiple European countries. The tree can be divided into two major branches resulting from the A23403G (D614G) mutation. The branch at the bottom is directly associated with the first recorded sequences from Wuhan, China and does not carry the D614G mutation. The Nigerian samples cluster with these early Chinese samples in the bottom branch of the tree. The branch on top is associated with sequences prevalent in Europe as demonstrated by reference sequences from Germany, France, Italy, Austria, Netherlands and UK. Ghanaian samples are about equally distributed between the top (European) and bottom branch of the tree. Senegalese samples cluster close with the French reference sample at the top of the tree.

**Fig. 1:**
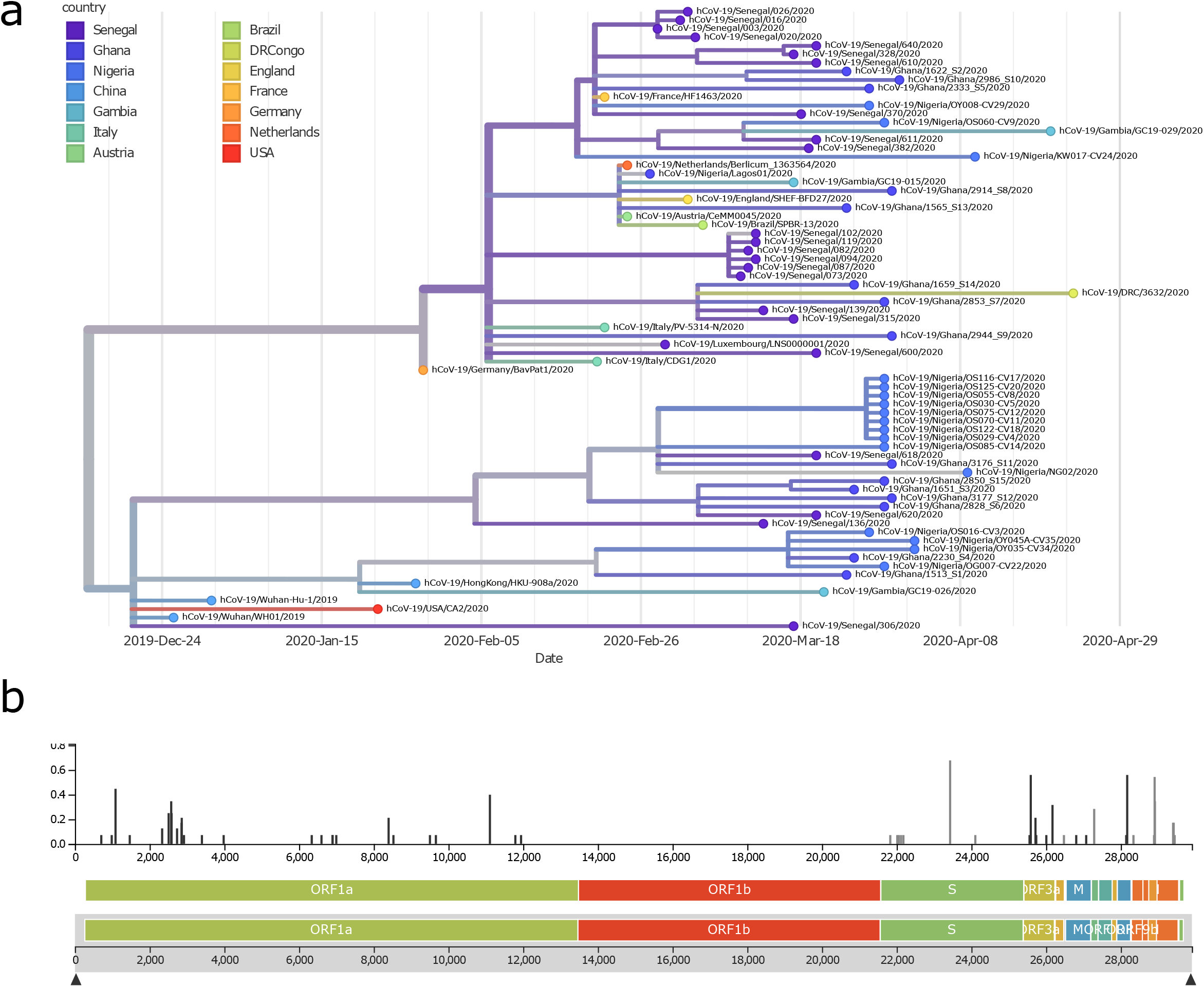
Phylogenetic tree revealing similarities of West-African viral sequences with Chinese and multiple European countries. (a) Nigerian samples cluster with the early Chinese samples within the bottom branch of the tree, Ghanaian samples are about equally distributed between the top (European) and bottom branch of the tree. Senegalese samples cluster closer with the French reference sample on the top of the tree. (b) Highest diversity is at the A23403G (D614G) mutation splitting the tree in the bottom (Chinese) and top (European) branch. This mutation was reported to increase infectivity.

The phylogenetic tree can be viewed interactively via the nextstrain.org framework under the URL: https://nextstrain.org/community/wwruck/wa

The split of the tree by the A23403G (D614G) mutation into two major branches corresponds to the highest diversity found at that location (Figure 1b). This mutation resides within the spike protein.

### Association with clades

We associated the West-African and reference samples via their characteristic mutations with clades according to the GISAID nomenclature. The phylogenetic tree in Figure 2 is coloured by these clades. The West-African samples are distributed over all clades suggesting introductions from China and European countries. However, each of the investigated countries has a specific pattern: most Senegalese samples have close similarity with the French reference, most Nigerian samples cluster in early Chinese-based clade S and Ghanaian samples are spread over all clades, the three Gambian samples are distributed over clades V, GR and GH. Within the clade S, there are putatively specific West-African mutations at the branches at C24370T and G22468T. Ghanaian samples predominate in the branch associated with the C24370T mutation. The branch determined by the mutation G22486T (Supplementary Figure 1) may reflect migration routes because in the nextstrain analysis of whole Africa there are also samples from Mali and Tunisia in this branch (https://nextstrain.org/ncov/africa?f_region=Africa, accessed August 14 2020). Two of the non-French-related Senegalese samples emanate from the C24370T and G22468T branches while the other (Senegal/136) has strong similarity with Spanish end-February samples from the early clade S (Supplementary Figure 2) pointing at multiple introductions to Senegal from France, Spain and African countries.

**Fig. 2:**
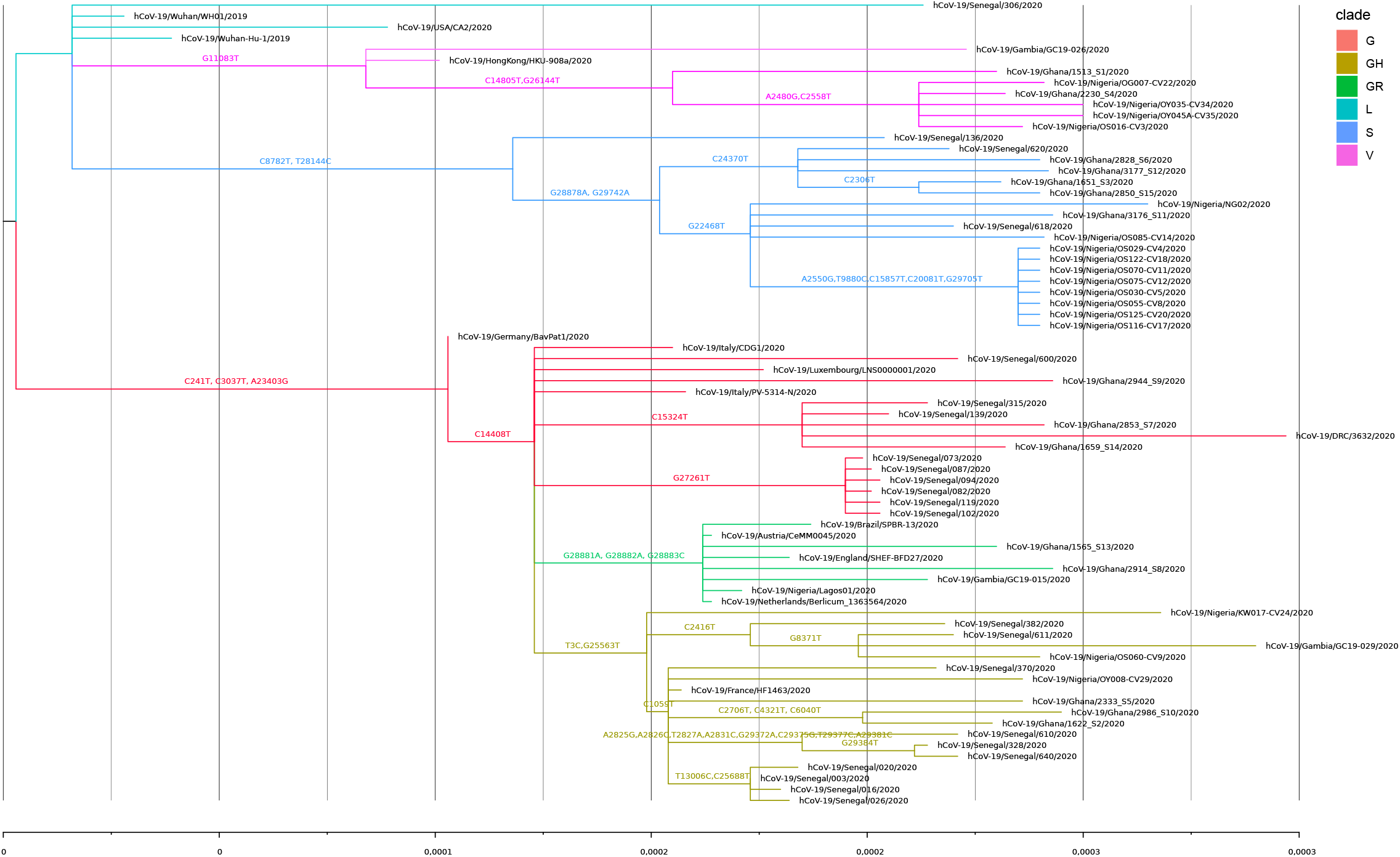
Phylogenetic tree colored by clades shows distribtution of West-African samples over all clades suggesting introductions from China and European countries. Patterns are country.specific, e.g. most Senegalese samples have close similarity with the French reference, most Nigerian samples cluster in early Chinese-based clade S and Ghanaian samples are spread over all clades. Within the clade S, there are putatively specific West-African mutations at the branches at C24370T and G22468T. G22486T may reflect migration routes because in the nextstrain analysis of whole Africa there are also Tunisian samples in this branch (https://nextstrain.org/ncov/africa?f_region=Africa, accessed Jun 26th, 2020). Two of the non-French related Senegalese samples come from these branches while the other (Senegal/136) has strong similarity with Spanish-end of February samples from the early clade S pointing at multiple introductions to Senegal from France, Spain and African countries.

### Timeline of clade distribution

In the temporal course of the clade distribution in Figure 3, the increased share of the Europe-associated G-clades becomes obvious. The G-clades harbor the putatively more infective D614G mutation (1). Surprisingly, the later Europe-associated G-clades (G, GH, GR) emerged before the earlier clades L, S and V in West African sequenced samples. This could be due to founder effects by introductions from France closely connected to Senegal and displaying a similar clade distribution and by migration and travel routes such as in the first registered Nigerian case infected in Italy (23). Furthermore, the China-based L-,V- and S-clade samples were obtained in mid-March, a time point within the Wuhan lockdown and when the epidemic in China was nearly totally over. Thus, the virus may have circulated in several countries before the first samples were sequenced. Surprisingly, the abundance of the S-clade is relatively high mainly due to the contribution from Nigeria and Ghana. However, without the S-clade distribution, the change in abundance resembles the global one with a delay of about 2-4 weeks.

**Fig. 3:**
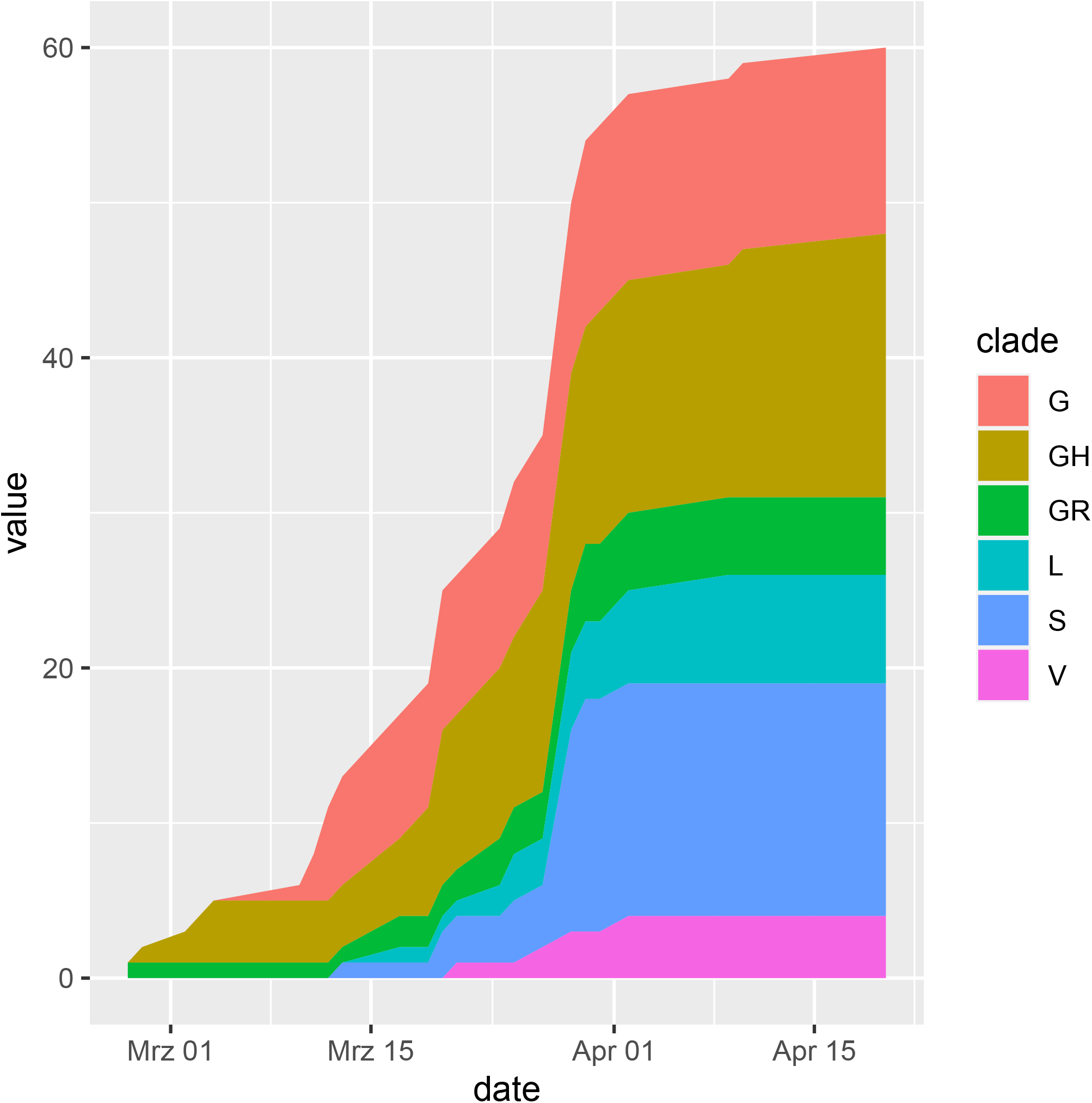
Temporal course of clade distribution confirms gaining of share of the Europe-associated G-clades harboring the putatively more infective D614G mutation. Interestingly the younger Europe-associated G-clades emerged earlier in West African sequenced samples. This could be due to founder effects by introductions from France closely connected to Senegal and displaying a similar clade distribution. Furthermore, the China-based L-,V- and S-clade samples start in mid-March a time when the epidemic in China was nearly totally suppressed. Thus, the virus may have circulated in several countries before the first samples were sequenced. Surprisingly, the abundance of the S-clade is relatively high mainly due to Nigeria and Ghana but without that exception the clade distribution resembles the global one with a delay of about 2-4 weeks.

### Country-specific patterns of clade distribution

Figure 4 shows that West African countries have acquired distinct patterns of China-and Europe-based clades. The first row contains the clade distribution charts of the West African countries investigated here whilst the second row contains charts of countries with comparable distributions. Nigeria has the highest percentage of the China-based early clades (L,S,V). Ghana has nearly equally distributed percentages of China and Europe-based clades (G,GH,GR) and in that sense has similarities with the German distribution. Senegal’s clade distribution resembles the one from France but includes also a few samples from the early China-based clades. There were only three sequences from Gambian, two from Europe-based clades GR and GH and one from China-based clade V. That pattern resembles the one from Italy when the clade G is substituted by the G-derived GH clade which however does not infer a connection to Italy but instead a similar combination of Chinese and European-related clades. Also the UK distribution in the last row has similarity with the Gambian distribution but as it includes also Chinese clades it also resembles the one from Ghana. The Dutch distribution which is quite similar to the German also resembles the clade distribution from Ghana. Last but not least, there are the quite distinct distribution from the US West and East Coast (California, CA and New York, NY). The Californian chart has similarity with the Nigerian because of the high percentage of Chinese-based clades while the chart from New York has a comparable high percentage of clade GH as the ones from France and Senegal.

**Fig. 4:**
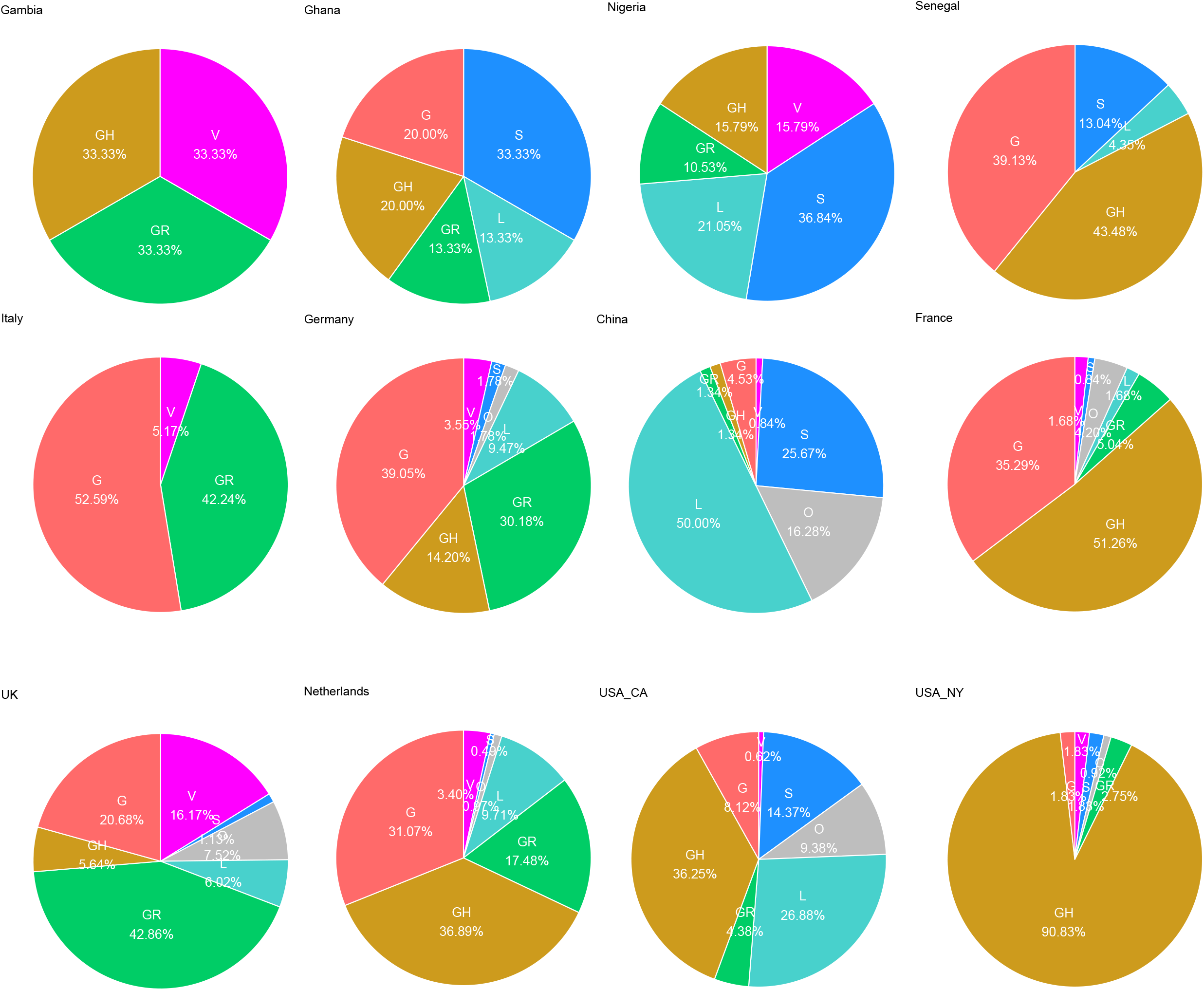
West African countries display distinct patterns of China-and Europe-based clades. Nigeria has the highest percentage of the China-based early clades (L,S,V) and Ghana has nearly equally distributed percentages of China and Europe-based clades (G,GH,GR). Senegal has a similar clade distribution as France but also a few samples from the early China-based clades. In Gambia there were only three sequences, two from Europe-based clades GR and GH and one from China-based clade V.

### Geographic distribution

The world map in Figure 5 reveals the distinct combinations of introduction of China-and Europe-based clades in West African countries. Nigeria has the highest percentage of the early clades (L,S,V) which were based in China but subsequently distributed to Europe and to the West Coast of USA. Ghana possesses nearly equally distributed percentages of the early clades and the Europe-based clades (G,GH,GR) comparable to the West Coast of USA. Senegal has a similar clade distribution like France and only few samples from the early China-based clades might be more comparable to the US East Coast.

**Fig. 5:**
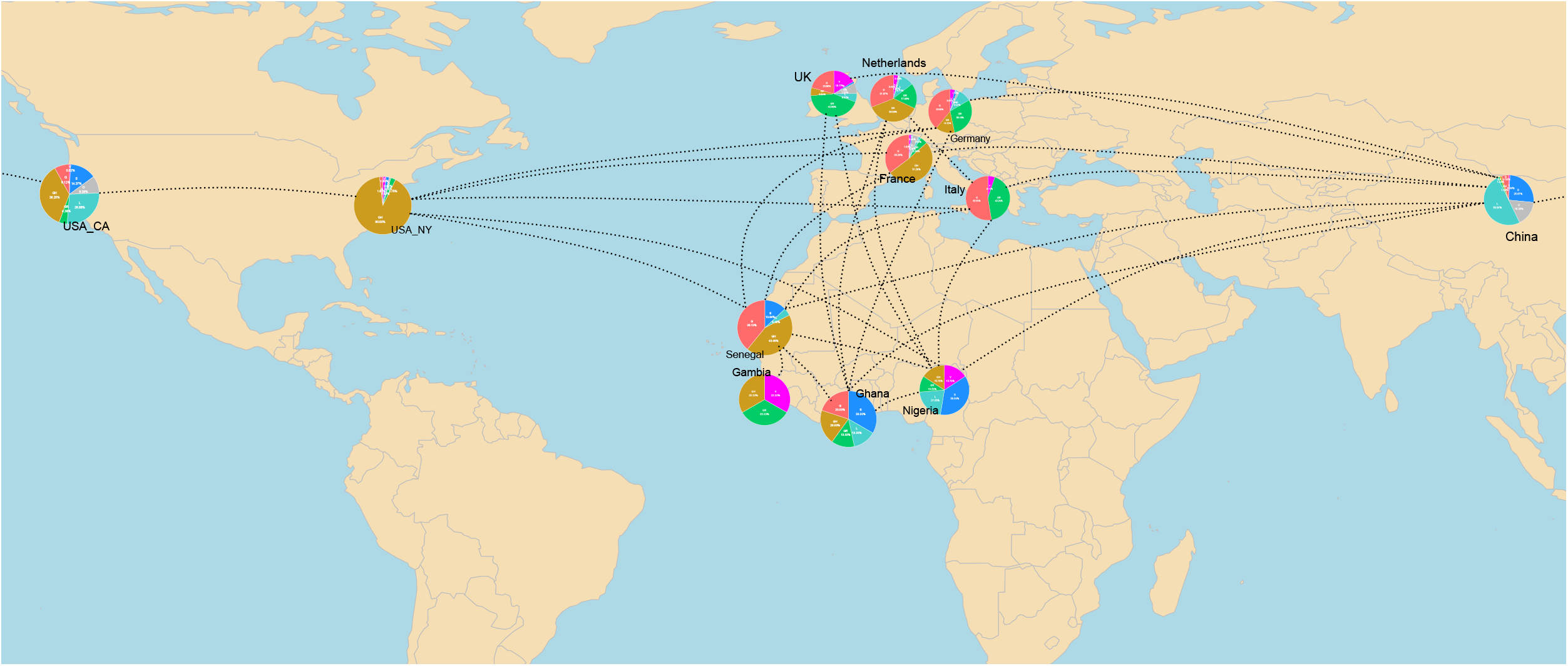
Geographic map reveals distinct patterns of introduction of China-and Europe-based clades in West African countries. Nigeria with the highest percentage of the China-based early clades (L,S,V) and Ghana with nearly equally distributed percentages of China and Europe-based clades (G,GH,GR) might be comparable with the US West Coast while Senegal with a similar clade distribution like France and few samples from the early China-based clades may be more comparable to the US East Coast. It will be interesting to observe if the later G clades replace the early clades in Nigeria and Ghana and if that correlates with the severity of the disease as was postulated for the US.

We set out to further explore the above-mentioned surprising observation (Figure 3) that in West Africa the early clades emerged after the later Europe-associated G-clades. Possible explanations could be (i) latent circulation of the early clades in West Africa or (ii) later introduction of the earlier clades. With the aim to find evidence for one of these alternatives, we looked into detail of the phylogeny of samples from the earlier clades. We picked two samples from the early clades: sample Senegal/136/2020 comes from a phylogenetic branch predominated by Spanish samples but also including samples from Asia and Latin America (suppl. Figure 2), several West African samples from Nigeria (dated March 29^th^, 2020), Ghana and Senegal in the phylogenetic branch in suppl. Figure 3 have a long latency time of about 2 months to the estimated common predecessor estimated on January 29^th^, 2020. Thus, there is evidence for a combination of both explanations : SARS-CoV-2 samples of the early clades may have circulated latently in West Africa since January 2020 but additionally there might have been introductions of the early clades from Europe and Asia or via maritime trade.

## Discussion

In this phylogenetic analysis of SARS-CoV-2 sequences from the West African countries Gambia, Ghana, Nigeria and Senegal we identified country-specific patterns of earlier (L, S, V) and later Europe-associated (G,GR, GH) clades. In Senegal and Gambia, the later Europe-associated clades were predominant, in Ghana earlier and later clades were more equally distributed and in Nigeria the earlier clades were the predominant samples downloaded from the GISAID database in June 2020. This would suggest multiple introductions mainly from Europe into Senegal and Gambia, from Europe and directly or indirectly via other Asian or European countries from China then to Ghana and Nigeria. The introductions from China to Nigeria and Ghana are in line with a study by Haider *et al*. in which both countries, but not Senegal and Gambia, appear in a table of estimations of SARS-CoV-2 transmission risk from China based on air traffic statistics (10). However, they are at low risk in the second quartile - with the fourth quartile having the highest risk. There was a lack of data for Senegal and Gambia therefore hinting to no or only low-level air traffic connection to China, thus suggesting a predominant introduction from Europe.

Against our expectations, we found that the later European-associated clades (G, GR, GH) emerged before the earlier Chinese-based clades (L, S, V) in the registered cases in the investigated West African countries. We propose the following hypothesis as an explanation to this surprising observation: the early clades were already circulating within the populations before the later European-associated clades were introduced. A higher disease severity of the later European clades might then be a possible explanation for their earlier detection. Intriguingly, most of the cases investigated in this study occurred within the time interval of the Wuhan lockdown between January 23^rd^ and April 8^th^ 2020. Thus, transmission of the early clades must have taken place very early or via intermediate countries or other Chinese provinces. Besides the later Europe-associated G-clades, the early clades were also circulating in Europe and the US West coast of USA, for example, the Senegal sample no. 136 from the early clade S has similarity with Spanish samples (suppl. Figure 2). Other explanations for the relatively long latency may be founder effects that by chance individuals infected with the later clades travelled to West Africa before individuals infected with the earlier clades – or slower means of transportation such as ships commuting between China, America, Europe and West Africa.

Based on previous reports (1), it might probably be that the later G clades will replace the early clades in Nigeria and Ghana. The question if that correlates with the severity of the disease still needs to be addressed, Brufsky infers it from the higher mortality at the East Coast of USA with predominantly D614G-carrying G–clades compared to the West Coast with the predominant early clades (2). Becerra-Flores et al. found significant correlations between the percentage of D614G and case-fatality on a country by country basis (24). However, others find evidence for higher transmissibility and also higher viral-load but no evidence for higher disease severity (1), (25), (26). A correlation of the mutation D614G associated with the G-clades and case fatality in the West African countries can only be identified at a marginal level of r=0.28 (Supplementary Table 1). The case fatality is fortunately rather low ranging from 0.6 in Ghana up to 3.2 in Gambia. Other factors such as climate, sunlight exposure (27) and associated Vitamin D (28), medical infrastructure and demographics might influence the etiopathology even more. There are also perspectives of decreased disease severity as Benedetti *et al*. argue that SARS-CoV-2 will mutate continuously and attenuate naturally to become endemic at a low mortality rate (29), as has been observed with earlier viruses (30).

The limitations of this study are the sample size, possible selection bias of the samples and the intrinsic incompleteness of the phylogenetic analysis which may lead to altered results when more samples are included. Nonetheless, this is the first study of its kind, the data and concept should form the basis for a more extensive analysis due to an increased number of sequenced samples.

In conclusion, in this phylogenetic analysis of SARS-CoV-2, we found distinct patterns of viral clades: the later Europe-associated G-clades are predominant in Senegal and Gambia, and combinations of the earlier (L, S, V) and later clades in Ghana and Nigeria. Intriguingly, the later clades emerged before the earlier clades which could simply be due to founder effects or due to latent circulation of the earlier clades. Only a marginal correlation of the G-clades in the West African countries can be associated with mortality which fortunately is at a rather low level therefore disproving fears that the pandemic would massively overwhelm the health systems in Africa. The rather young population and the climate might be factors favoring this low fatality rate in comparison to Western countries but nevertheless a cautious balance between health protection and economics might prevent future disastrous outbreaks.

## Author Contributions

JA and WW came up with the concept of the study. WW and JA wrote the manuscript. WW analysed and WW and JA interpreted the data. JA supervised the work.

## Acknowledgments

We gratefully acknowledge all authors from the originating laboratories and from the submitting laboratories of the sequences from GISAID and GenBank which are listed in Supplementary Table 2. James Adjaye acknowledges financial support from the medical faculty of Heinrich Heine University-Düsseldorf, Germany.

## Supplementary Material

**Supplementary Table 1: Case fatality and percentage of mutation D614G in West African countries**

**Supplementary Table 2: Acknowledgement table of sequence samples from the GISAID database**

**Supplementary Figure 1: Detailed phylogenetic analysis of the Senegal/618 and several Nigerian samples point at introduction through travel or migration routes via Tunisia, Egypt and Mali.**

**Supplementary Figure 2: Detailed phylogenetic analysis of the Senegal/136 sample suggests introduction from Spain.**

**Supplementary Figure 3: Detailed phylogenetic analysis of Nigerian, Ghanian and Senegalese samples points at long latent circulation of early clades of SARS-CoV-2 in these countries between end of January until end of March 2020.**

