## Supplementary figures and images for "Transmission of SARS-COV-2 from China to Europe and West Africa: a detailed phylogenetic analysis"

### Supplementary Figure 1

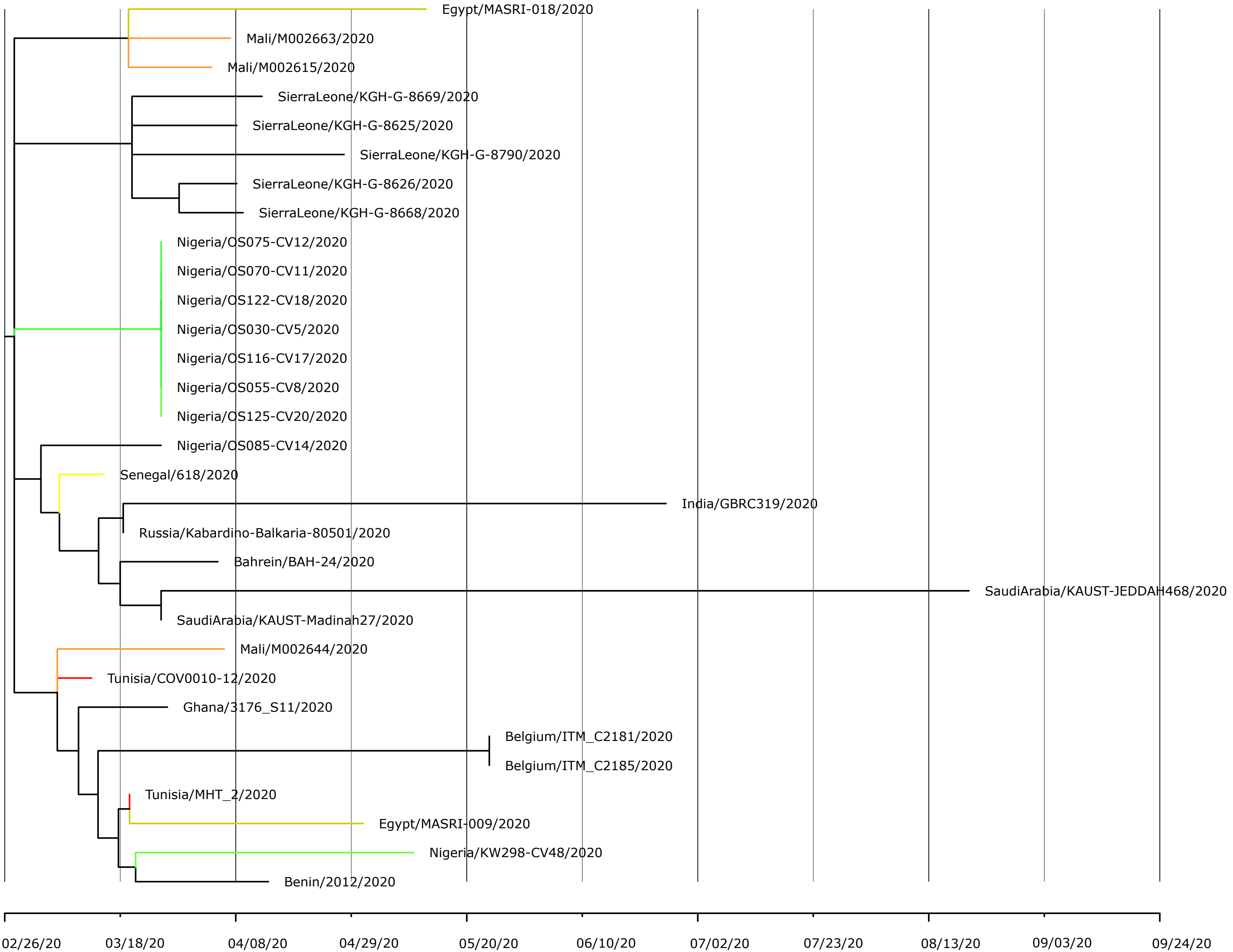

### Supplementary Figure 2

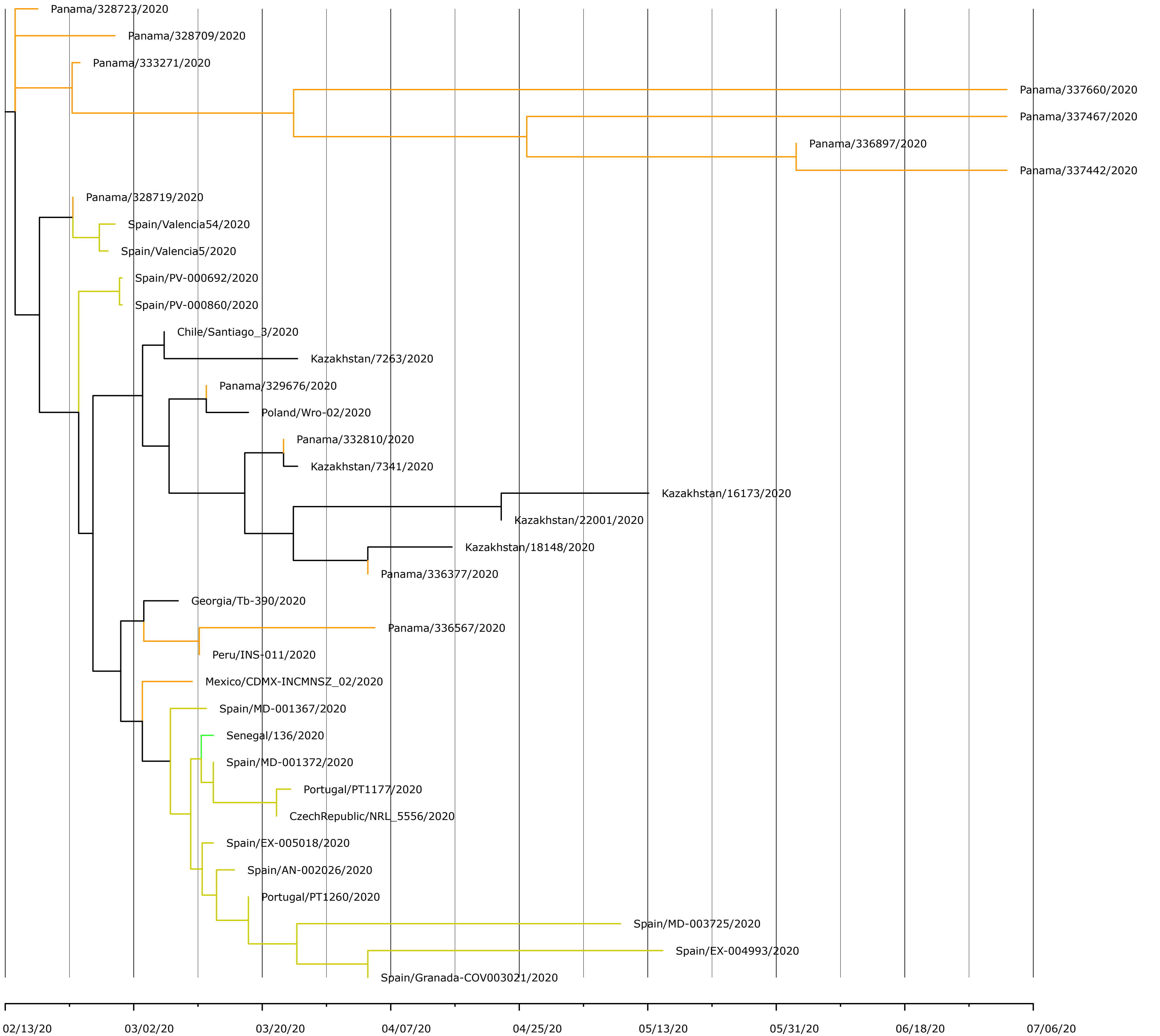

### Supplementary Figure 3

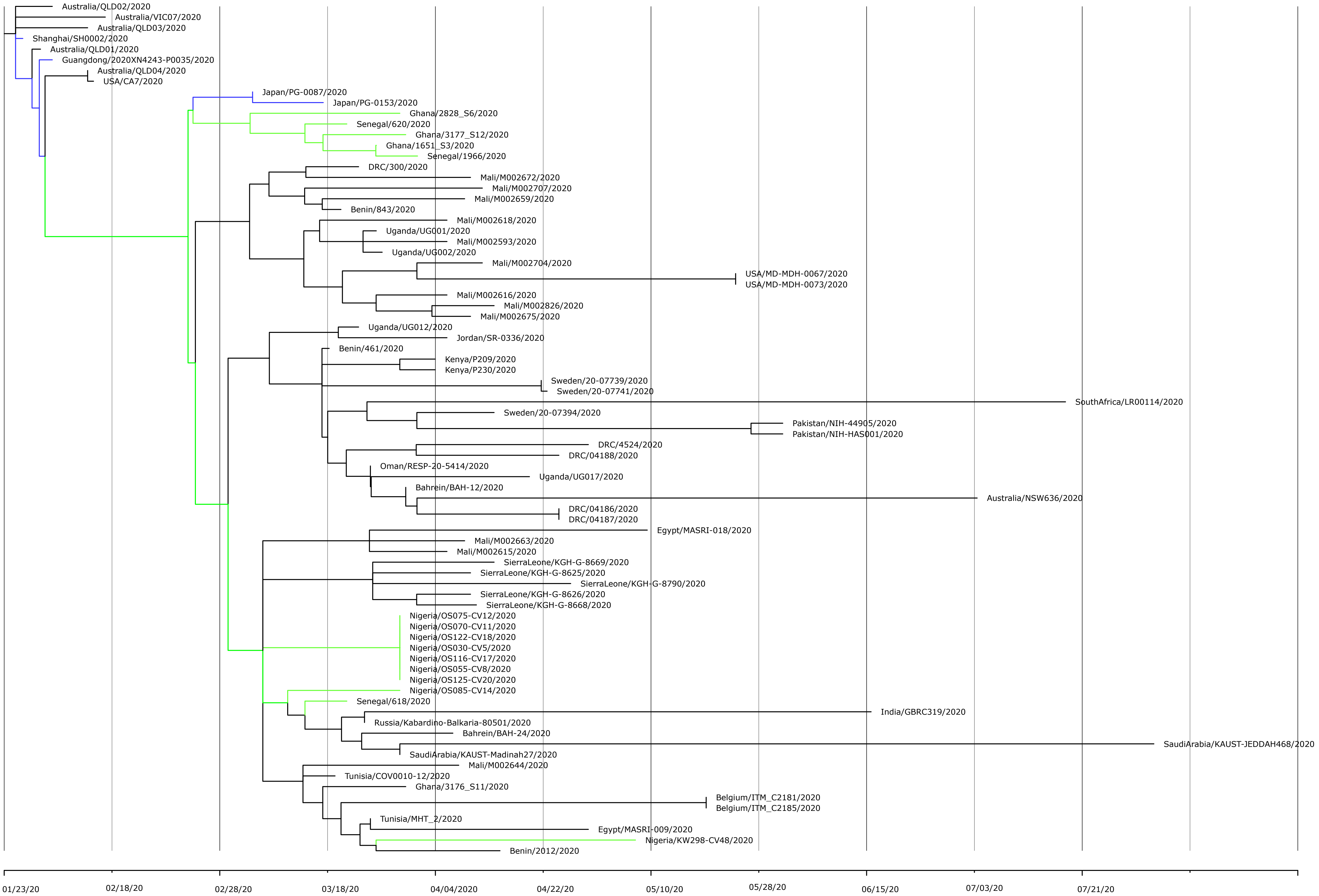
