## Supplementary Table 1 for "Transmission of SARS-COV-2 from China to Europe and West Africa: a detailed phylogenetic analysis"

**Supplementary Table 1: Case fatality and percentage of mutation D614G in West African countries**

| country | deaths | cases | case fatality (%) | D614G (%) |
| --- | --- | --- | --- | --- |
| Senegal | 272 | 13013 | 2.09 | 82.61 |
| Gambia | 87 | 2685 | 3.24 | 66.67 |
| Ghana | 261 | 43505 | 0.60 | 53.33 |
| Nigeria | 1002 | 52227 | 1.92 | 26.32 |
|  |  |  | Pearson correlation  r(% D614G, % case fatality): | 0.28 |

(Retrieved from the Johns Hopkins University corona map <https://coronavirus.jhu.edu/map.html>, 08/24/20)
