## Supplementary Table 2 for "Transmission of SARS-COV-2 from China to Europe and West Africa: a detailed phylogenetic analysis"

We gratefully acknowledge the following Authors from the Originating laboratories responsible for obtaining the specimens and the

Submitting laboratories where genetic sequence data were generated and shared via the GISAID Initiative, on which this research is based.

| Virus name | Accession No. | Collected | Originating laboratory | Submitting laboratory | authors |
| --- | --- | --- | --- | --- | --- |
| Austria/CeMM0045/2020 | EPI_ISL_437932 | 24.02.2020 | Institut für Virologie am Department für Hygiene, Mikrobiologie und Public Health | Bergthaler laboratory, CeMM Research Center for Molecular Medicine of the Austrian Academy of Sciences | Alexandra Popa et al  (1) |
| Brazil/SPBR-13/2020 | EPI_ISL_416035 | 05.03.2020 | National Influenza Center - Instituto Adolfo Lutz | Instituto Adolfo Lutz, Interdiciplinary Procedures Center, Strategic Laboratory | Claudio Tavares Sacchi et al  (2) |
| DRC/3632/2020 | EPI_ISL_447607 | 23.04.2020 | Viral Respiratory Lab, National Institute for Biomedical Research (INRB) | Pathogen Sequencing Lab, National Institute for Biomedical Research (INRB) | Placide Mbala-Kingebeni et al  <https://virological.org/t/phylogenetic-analysis-of-sars-cov-2-in-drc/528>  [accessed 08/18/20] |
| England/SHEF-BFD27/2020 | EPI_ISL_416737 | 03.03.2020 | Virology Department, Sheffield Teaching Hospitals NHS Foundation Trust | Department of Infection, Immunity and Cardiovascular Disease, The Florey Institute, The Medical School, University of Sheffield | Thushan de Silva et al  <https://virological.org/t/preliminary-analysis-of-sars-cov-2-importation-establishment-of-uk-transmission-lineages/507>  [accessed 08/18/20] |
| France/HF1463/2020 | EPI_ISL_429968 | 21.02.2020 | Centre Hospitalier Compiègne Laboratoire de Biologie | National Reference Center for Viruses of Respiratory Infections, Institut Pasteur, Paris | Mélanie Albert et al  (3) |
| Gambia/GC19-015/2020 | EPI_ISL_428855 | 17.03.2020 | MRCG at LSHTM Geomics lab | MRCG at LSHTM Genomics lab | Sesay et al et al (4) |
| Gambia/GC19-026/2020 | EPI_ISL_428856 | 21.03.2020 | MRCG at LSHTM Genomics Lab | MRCG at LSHTM Genomics lab | Sesay et al et al (4) |
| Gambia/GC19-029/2020 | EPI_ISL_428857 | 20.04.2020 | MRCG at LSHTM Genomics lab | MRCG at LSHTM Genomics lab | Sesay et al et al (4) |
| Germany/BavPat1/2020 | EPI_ISL_406862 | 28.01.2020 | Charité Universitätsmedizin Berlin, Institute of Virology; Institut für Mikrobiologie der Bundeswehr, Munich | Charité Universitätsmedizin Berlin, Institute of Virology | Victor M Corman et al (5) |
| Ghana/1513_S1/2020 | EPI_ISL_422382 | 24.03.2020 | NMIMR, Department of Virology | WACCBIP, University of Ghana | Joyce M. Ngoi et al |
| Ghana/1565_S13/2020 | EPI_ISL_422404 | 24.03.2020 | NMIMR, Department of Virology | WACCBIP, University of Ghana | Joyce M. Ngoi et al |
| Ghana/1622_S2/2020 | EPI_ISL_422384 | 24.03.2020 | NMIMR, Department of Virology | WACCBIP, University of Ghana | Joyce M. Ngoi et al |
| Ghana/1651_S3/2020 | EPI_ISL_422387 | 25.03.2020 | NMIMR, Department of Virology | WACCBIP, University of Ghana | Joyce M. Ngoi et al |
| Ghana/1659_S14/2020 | EPI_ISL_422405 | 25.03.2020 | NMIMR, Department of Virology | WACCBIP, University of Ghana | Joyce M. Ngoi et al |
| Ghana/2230_S4/2020 | EPI_ISL_422390 | 25.03.2020 | NMIMR, Department of Virology | WACCBIP, University of Ghana | Joyce M. Ngoi et al |
| Ghana/2333_S5/2020 | EPI_ISL_422394 | 27.03.2020 | NMIMR, Department of Virology | WACCBIP, University of Ghana | Joyce M. Ngoi et al |
| Ghana/2828_S6/2020 | EPI_ISL_422397 | 29.03.2020 | NMIMR, Department of Virology | WACCBIP, University of Ghana | Joyce M. Ngoi et al |
| Ghana/2850_S15/2020 | EPI_ISL_422406 | 29.03.2020 | NMIMR, Department of Virology | WACCBIP, University of Ghana | Joyce M. Ngoi et al |
| Ghana/2853_S7/2020 | EPI_ISL_422398 | 29.03.2020 | NMIMR, Department of Virology | WACCBIP, University of Ghana | Joyce M. Ngoi et al |
| Ghana/2914_S8/2020 | EPI_ISL_422399 | 30.03.2020 | NMIMR, Department of Virology | WACCBIP, University of Ghana | Joyce M. Ngoi et al |
| Ghana/2944_S9/2020 | EPI_ISL_422400 | 30.03.2020 | NMIMR, Department of Virology | WACCBIP, University of Ghana | Joyce M. Ngoi et al |
| Ghana/2986_S10/2020 | EPI_ISL_422401 | 31.03.2020 | NMIMR, Department of Virology | WACCBIP, University of Ghana | Joyce M. Ngoi et al |
| Ghana/3176_S11/2020 | EPI_ISL_422402 | 30.03.2020 | NMIMR, Department of Virology | WACCBIP, University of Ghana | Joyce M. Ngoi et al |
| Ghana/3177_S12/2020 | EPI_ISL_422403 | 30.03.2020 | NMIMR, Department of Virology | WACCBIP, University of Ghana | Joyce M. Ngoi et al |
| HongKong/HKU-908a/2020 | EPI_ISL_434569 | 27.01.2020 | unknown | Microbiology | To et al |
| Italy/CDG1/2020 | EPI_ISL_412973 | 20.02.2020 | Department of Infectious Diseases, Istituto Superiore di Sanità, Roma , Italy | Virology Laboratory, Scientific Department, Army Medical Center | Paola Stefanelli et al (5) |
| Italy/PV-5314-N/2020 | EPI_ISL_451307 | 21.02.2020 | Molecular Virology Unit, Fondazione IRCCS Policlinico San Matteo , Pavia | Laboratory of Virology, INMI Lazzaro Spallanzani IRCCS | Fausto Baldanti et al |
| Luxembourg/LNS0000001/2020 | EPI_ISL_419562 | 29.02.2020 | Laboratoire National de Santé, Microbiology, Virology | Laboratoire National de Santé, Microbiology, Epidemiology and Microbial Genomics | Anke Wienecke-Baldacchino et al |
| Netherlands/Berlicum_1363564/2020 | EPI_ISL_413565 | 24.02.2020 | Foundation Pamm | Erasmus Medical Center | David Nieuwenhuijse et al (6) |
| Nigeria/KW017-CV24/2020 | EPI_ISL_455362 | 10.04.2020 | Nigeria Centre for Disease Control (NCDC) | African Centre of Excellence for Genomics of Infectious Diseases (ACEGID), Redeemer's University, Ede, Osun State, Nigeria | Oluniyi P.E. et al  [https://virological.org/t/sars-cov-2-genomes-from-nigeria-reveal-community-transmission-multiple-virus-lineages-and-spike-protein-mutation-associated-with-higher-transmission-and-pathogenicity/494 2020-05-29](https://virological.org/t/sars-cov-2-genomes-from-nigeria-reveal-community-transmission-multiple-virus-lineages-and-spike-protein-mutation-associated-with-higher-transmission-and-pathogenicity/494%202020-05-29)  [accessed 08/18/20] |
| Nigeria/Lagos01/2020 | EPI_ISL_413550 | 27.02.2020 | Centre for Human and Zoonotic Virology (CHAZVY), College of Medicine University of Lagos/Lagos University Teaching Hospital (LUTH), part of the Laboratory Network of the Nigeria Centre for Disease Control (NCDC) | African Centre of Excellence for Genomics of Infectious Diseases (ACEGID), Redeemer's University, Ede, Osun State, Nigeria | Oluniyi P.E. et al |
| Nigeria/NG57752/2020 | EPI_ISL_462992 | 2020-03 | unknown | Director General | Saibu et al |
| Nigeria/OG007-CV22/2020 | EPI_ISL_455412 | 29.03.2020 | Nigeria Centre for Disease Control (NCDC) | African Centre of Excellence for Genomics of Infectious Diseases (ACEGID), Redeemer's University, Ede, Osun State, Nigeria | Oluniyi P.E. et al |
| Nigeria/OS016-CV3/2020 | EPI_ISL_455413 | 27.03.2020 | Nigeria Centre for Disease Control (NCDC) | African Centre of Excellence for Genomics of Infectious Diseases (ACEGID), Redeemer's University, Ede, Osun State, Nigeria | Oluniyi P.E. et al |
| Nigeria/OS029-CV4/2020 | EPI_ISL_455414 | 29.03.2020 | Nigeria Centre for Disease Control (NCDC) | African Centre of Excellence for Genomics of Infectious Diseases (ACEGID), Redeemer's University, Ede, Osun State, Nigeria | Oluniyi P.E. et al |
| Nigeria/OS030-CV5/2020 | EPI_ISL_455415 | 29.03.2020 | Nigeria Centre for Disease Control (NCDC) | African Centre of Excellence for Genomics of Infectious Diseases (ACEGID), Redeemer's University, Ede, Osun State, Nigeria | Oluniyi P.E. et al |
| Nigeria/OS055-CV8/2020 | EPI_ISL_455418 | 29.03.2020 | Nigeria Centre for Disease Control (NCDC) | African Centre of Excellence for Genomics of Infectious Diseases (ACEGID), Redeemer's University, Ede, Osun State, Nigeria | Oluniyi P.E. et al |
| Nigeria/OS060-CV9/2020 | EPI_ISL_455419 | 29.03.2020 | Nigeria Centre for Disease Control (NCDC) | African Centre of Excellence for Genomics of Infectious Diseases (ACEGID), Redeemer's University, Ede, Osun State, Nigeria | Oluniyi P.E. et al |
| Nigeria/OS070-CV11/2020 | EPI_ISL_455422 | 29.03.2020 | Nigeria Centre for Disease Control | African Centre of Excellence for Genomics of Infectious Diseases (ACEGID), Redeemer's University, Ede, Osun State, Nigeria | Oluniyi P.E. et al |
| Nigeria/OS075-CV12/2020 | EPI_ISL_455423 | 29.03.2020 | Nigeria Centre for Disease Control (NCDC) | African Centre of Excellence for Genomics of Infectious Diseases (ACEGID), Redeemer's University, Ede, Osun State, Nigeria | Oluniyi P.E. et al |
| Nigeria/OS085-CV14/2020 | EPI_ISL_455424 | 29.03.2020 | Nigeria Centre for Disease Control (NCDC) | African Centre of Excellence for Genomics of Infectious Diseases (ACEGID), Redeemer's University, Ede, Osun State, Nigeria | Oluniyi P.E. et al |
| Nigeria/OS116-CV17/2020 | EPI_ISL_455425 | 29.03.2020 | Nigeria Centre for Disease Control (NCDC) | African Centre of Excellence for Genomics of Infectious Diseases (ACEGID), Redeemer's University, Ede, Osun State, Nigeria | Oluniyi P.E. et al |
| Nigeria/OS122-CV18/2020 | EPI_ISL_455426 | 29.03.2020 | Nigeria Centre for Disease Control | African Centre of Excellence for Genomics of Infectious Diseases (ACEGID), Redeemer's University, Ede, Osun State, Nigeria | Oluniyi P.E. et al |
| Nigeria/OS125-CV20/2020 | EPI_ISL_455427 | 29.03.2020 | Nigeria Centre for Disease Control (NCDC) | African Centre of Excellence for Genomics of Infectious Diseases (ACEGID), Redeemer's University, Ede, Osun State, Nigeria | Oluniyi P.E. et al |
| Nigeria/OY008-CV29/2020 | EPI_ISL_455429 | 27.03.2020 | Nigeria Centre for Disease Control (NCDC) | African Centre of Excellence for Genomics of Infectious Diseases (ACEGID), Redeemer's University, Ede, Osun State, Nigeria | Oluniyi P.E. et al |
| Nigeria/OY035-CV34/2020 | EPI_ISL_455430 | 02.04.2020 | Nigeria Centre for Disease Control (NCDC) | African Centre of Excellence for Genomics of Infectious Diseases (ACEGID), Redeemer's University, Ede, Osun State, Nigeria | Oluniyi P.E. et al |
| Nigeria/OY045A-CV35/2020 | EPI_ISL_455431 | 02.04.2020 | Nigeria Centre for Disease Control (NCDC) | African Centre of Excellence for Genomics of Infectious Diseases (ACEGID), Redeemer's University, Ede, Osun State, Nigeria | Oluniyi P.E. et al |
| Senegal/003/2020 | EPI_ISL_418206 | 28.02.2020 | Institut Pasteur Dakar | Institut Pasteur de Dakar | Ndongo Dia et al |
| Senegal/016/2020 | EPI_ISL_418207 | 02.03.2020 | Institut Pasteur Dakar | Institut Pasteur de Dakar | Ndongo Dia et al |
| Senegal/020/2020 | EPI_ISL_418208 | 04.03.2020 | Institut Pasteur Dakar | Institut Pasteur de Dakar | Ndongo Dia et al |
| Senegal/026/2020 | EPI_ISL_418209 | 03.03.2020 | Institut Pasteur Dakar | Institut Pasteur de Dakar | Ndongo Dia et al |
| Senegal/073/2020 | EPI_ISL_418210 | 10.03.2020 | Institut Pasteur Dakar | Institut Pasteur de Dakar | Ndongo Dia et al |
| Senegal/082/2020 | EPI_ISL_418211 | 11.03.2020 | Institut Pasteur Dakar | Institut Pasteur de Dakar | Ndongo Dia et al |
| Senegal/087/2020 | EPI_ISL_418212 | 11.03.2020 | Institut Pasteur Dakar | Institut Pasteur de Dakar | Ndongo Dia et al |
| Senegal/094/2020 | EPI_ISL_418213 | 12.03.2020 | Institut Pasteur Dakar | Institut Pasteur de Dakar | Ndongo Dia et al |
| Senegal/102/2020 | EPI_ISL_418214 | 12.03.2020 | Institut Pasteur Dakar | Institut Pasteur de Dakar | Ndongo Dia et al |
| Senegal/119/2020 | EPI_ISL_418215 | 12.03.2020 | Instirut Pasteur Dakar | Institut Pasteur de Dakar | Ndongo Dia et al |
| Senegal/136/2020 | EPI_ISL_418216 | 13.03.2020 | Institut Pasteur Dakar | Institut Pasteur de Dakar | Ndongo Dia et al |
| Senegal/139/2020 | EPI_ISL_418217 | 13.03.2020 | Institut Pasteur Dakar | Institut Pasteur de Dakar | Ndongo Dia et al |
| Senegal/306/2020 | EPI_ISL_420069 | 17.03.2020 | Institut Pasteur Dakar | Institut Pasteur de Dakar | Ndongo Dia et al |
| Senegal/315/2020 | EPI_ISL_420070 | 17.03.2020 | Institut Pasteur Dakar | Institut Pasteur de Dakar | Ndongo Dia et al |
| Senegal/328/2020 | EPI_ISL_420071 | 17.03.2020 | Institut Pasteur Dakar | Institut Pasteur de Dakar | Ndongo Dia et al |
| Senegal/370/2020 | EPI_ISL_420072 | 18.03.2020 | Institut Pasteur Dakar | Institut Pasteur de Dakar | Ndongo Dia et al |
| Senegal/382/2020 | EPI_ISL_420073 | 19.03.2020 | Institut Pasteur Dakar | Institut Pasteur de Dakar | Ndongo Dia et al |
| Senegal/600/2020 | EPI_ISL_420074 | 20.03.2020 | Institut Pasteur Dakar | Institut Pasteur de Dakar | Ndongo Dia et al |
| Senegal/610/2020 | EPI_ISL_420075 | 20.03.2020 | Institut pasteur Dakar | Institut Pasteur de Dakar | Ndongo Dia et al |
| Senegal/611/2020 | EPI_ISL_420076 | 20.03.2020 | Institut Pasteur Dakar | Institut Pasteur de Dakar | Ndongo Dia et al |
| Senegal/618/2020 | EPI_ISL_420077 | 20.03.2020 | Institut Pasteur Dakar | Institut Pasteur de Dakar | Ndongo Dia et al |
| Senegal/620/2020 | EPI_ISL_420078 | 20.03.2020 | Institut Pasteur Dakar | Institut Pasteur de Dakar | Ndongo Dia et al |
| Senegal/640/2020 | EPI_ISL_420079 | 20.03.2020 | Institut Pasteur Dakar | Institut Pasteur de Dakar | Ndongo Dia et al |
| USA/CA2/2020 | EPI_ISL_406036 | 22.01.2020 | California Department of Public Health | Pathogen Discovery, Respiratory Viruses Branch, Division of Viral Diseases, Centers for Disease Control and Prevention | Anna Uehara et al (7) |
| Wuhan/Hu-1/2019 | EPI_ISL_402125 | 26.12.2019 | unknown | National Institute for Communicable Disease Control and Prevention (ICDC) Chinese Center for Disease Control and Prevention (China CDC) | Zhang et al (8) |
| Wuhan/WH01/2019 | EPI_ISL_406798 | 26.12.2019 | General Hospital of Central Theater Command of People's Liberation Army of China | BGI & Institute of Microbiology, Chinese Academy of Sciences & Shandong First Medical University & Shandong Academy of Medical Sciences & General Hospital of Central Theater Command of People's Liberation Army of China | Weijun Chen et al (9) |
